# AI-Generated Virtual Libraries Could Help Uncover RNA-Specific Regions of Chemical Space

**DOI:** 10.1101/2022.02.05.479230

**Authors:** Ziqiao Xu, Aaron T. Frank

## Abstract

RNAs can recognize small-molecule ligands. However, the extent to which the molecules that they recognize differ from those recognized by proteins remains an open question. Cheminformatics analysis of experimentally validated RNA binders strongly suggests that RNA binders occupy a specific region of chemical space. However, less than 100 validated small molecule ligands are currently known. Here, we demonstrate how structure-based approaches could be used to navigate vast regions of the chemical space specific to ligand binding sites in five highly-structured RNAs. Our method involves using generative-AI to design target- and site-specific virtual libraries and then analyzing them using similar cheminformatics approaches as those used to assess experimentally validated RNA binders. Despite employing a completely orthogonal strategy, our results essentially reproduce the trends observed by analyzing the experimentally validated RNA binders. Large-scale generation of target and site-specific libraries may therefore prove to be helpful in simultaneously mapping the regions of chemical space unique to RNA and generating libraries that could be mined to identify novel RNA binders.

**TOC IMAGE:** 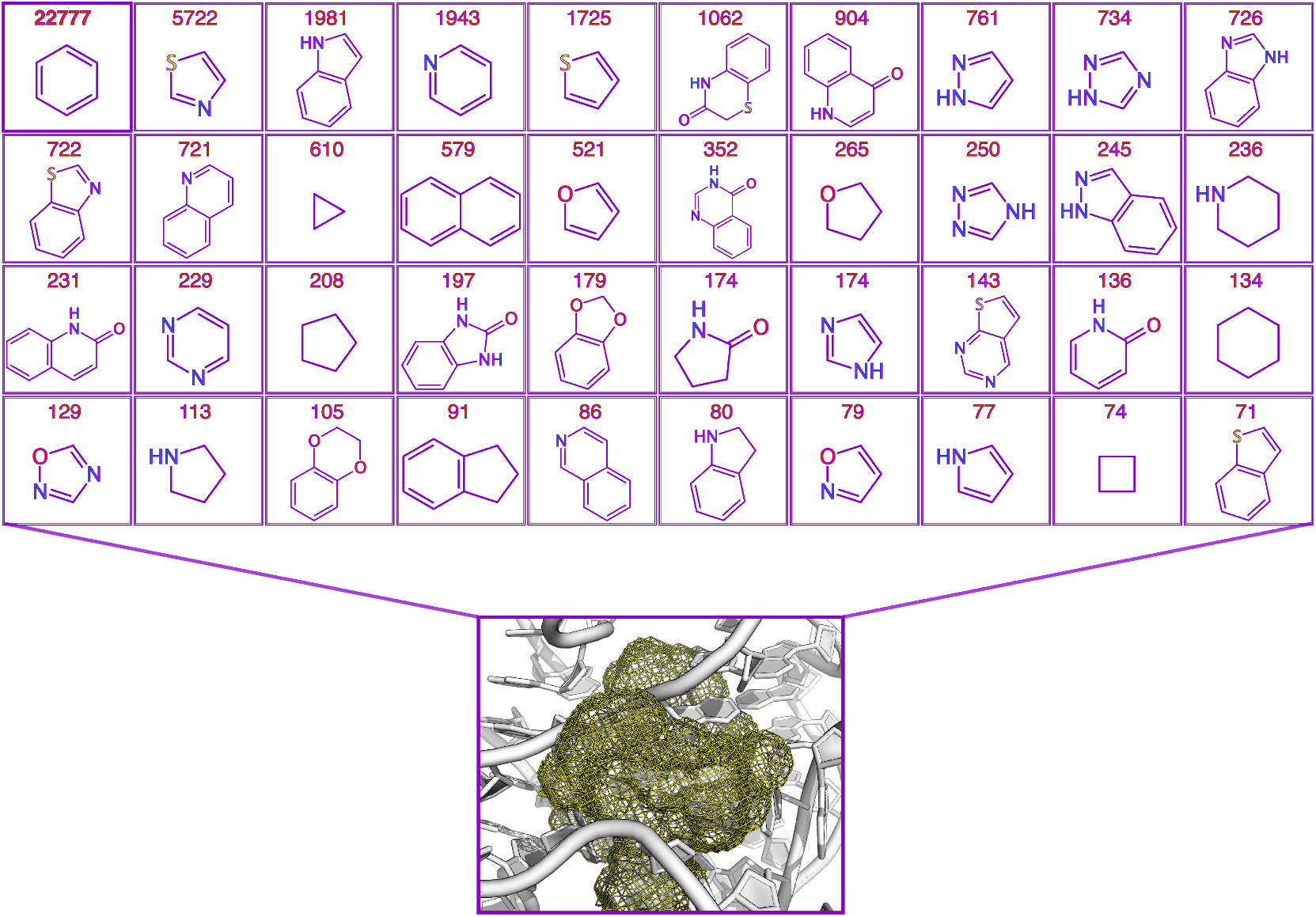

## INTRODUCTION

Ever since the discovery that some antibiotics directly target RNA elements within the ribosome,^1–3^ RNAs have been considered potential drug targets. The recent discovery and approval of the exon-7 targeting drug (risdiplam),^4,5^ which corrects SMN2 exon-7 mis-splicing in patients suffering from SMA, has elevated RNAs to the status of a bona vide drug targets. Also, the discovery of riboswitch targeting drug-like molecule (ribocil) with antibacterial activity against bacteria harboring the FMN riboswitch provides further support that identifying RNA-targeting some molecules is a viable drug development strategy.^6^ Further, the identification of antifungal molecules (by Pyle and coworkers) that target the highly structured group II intron and inhibit splicing,^7^ along with other recent discoveries,^8–12^ provides additional validation of the viability of RNAs, particularly highly structured RNAs, as drug targets.

Highly structured RNA can recognize small-molecule ligands that bind to cavities along its surface. Such small molecules can, in turn, modulate the dynamics of these RNAs and, in so doing, their function. Identifying or designing molecules that bind to an RNA is the first step in designing RNA-targeting ligands, serving as a starting point for drug development. Compared to proteins, however, we know little about which regions of chemical space contain small molecules that are likely to bind to RNA with high selective and specificity. Such rich information could prove helpful in guiding the exploration of chemical space in search of bioactive RNA-targeting small molecules. The answer to this question, namely, what are the regions of chemical-space that contain RNA-specific molecules, is beginning to trickle in. For example, recent work by Hargrove and coworkers indicate that RNA binders exhibit distributions along with key physicochemical properties that are distinct from FDA-approved protein targeting small molecules.^13,14^ Moreover, their analysis suggests that RNA binding ligands have a propensity for being rod-like. As additional examples of RNA targeting molecules become known,^15^ a complete picture of the chemical building blocks required to design RNA-specific molecules is likely to emerge.

In this study, we explored the utility of computer-aided structure-based methods in building up an intuition about the region(s) of chemical space that might be specific to RNA. Theoretical techniques, based on biophysical modeling (that explicitly considered the interactions with potential ligands), hold tremendous promise in mapping the regions of chemical space that are likely to contain RNA-targeting compounds because one can execute this mapping rapidly and at low cost. Therefore, in this study, we utilize a virtual library generation strategy to generate target/site-specific virtual libraries for five highly-structured RNAs. We first provide computational evidence that the libraries are target/site-specific, and then we analyze their collective properties. We found that the distributions of several physicochemical properties across our library overlap with the corresponding distribution across R-BIND, a curated database of known bioactive, RNA-targeting small molecules. Furthermore, consistent with previous analysis of the R-BIND molecules, those in our library also tended to be rod-like, though this trend was target/site-dependent. Finally, we found that frequently observed ring systems in our libraries were prevalent in the database of recently identified small fragments that target various elements in the SARS-CoV-2 RNA genome. Our results indicate that one could use existing structure-based methods to efficiently explore chemical space to predict the chemical building block of RNA-targeting ligands.

## METHODS

### RNA-ligand systems

To explore the utility of structure-guided techniques to explore chemical space specific to individual RNA-ligand cavities, we compile a set of five RNA-ligand systems (Table 1). These were chosen because they each are known to bind to small-molecule ligands, and high-resolution structures of these RNAs in complex with ligands are available in the PDB.

**Table 1:**
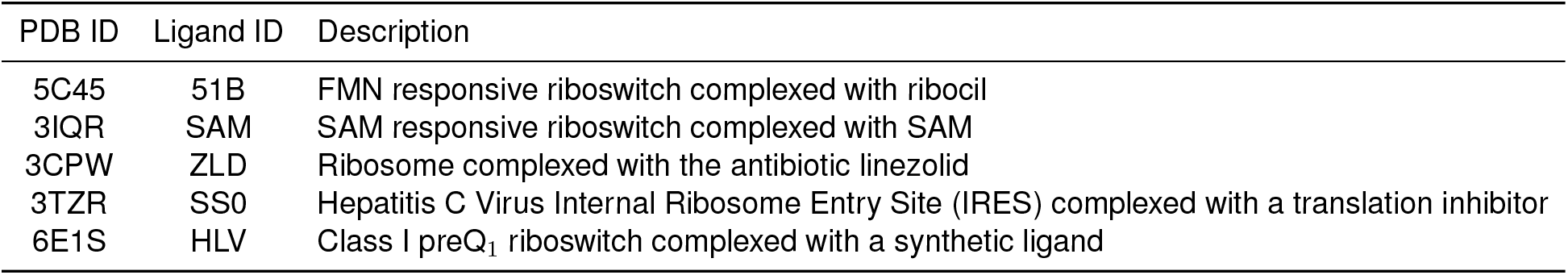
RNA targets used in this study.

### AI-guided chemical space exploration

To explore the regions of chemical space specific to the ligand-binding cavity in each of the five RNAs we examined in this study, we used the AI-assisted sample-and-dock approach.^16^ Briefly, the sample-and-dock framework uses generative AI and docking to iterative sampling points (molecules) in the neighborhood of an initial scaffold. These molecules are then docked into the cavity of the target, and the molecule with the lowest docking score is identified and used to seed the next sample-and-dock cycle. At the end of a fixed set of cycles, a library of molecules is compiled that is specific to the targeted cavity. Here we used the sample-and-dock framework implemented in the tool SampleDock. For each RNA, SampleDock was executed for 24 hours, resulting in ~3000 design cycles. The molecules with Z-scores of ≤ −1.0 from the docking score distributions were selected and used as the final library (see Table 1).

### Analysis of system-specific libraries

Mirroring work by Hargrove and coworkers,^14,17^ we calculated several properties from the molecules in our libraries. First, to assess the physicochemical properties of the molecules, we computed 18 of their cheminformatics parameters (see Table S1).^18^ Second, to characterize the distribution of shapes in our libraries, we calculated the principal moments of inertia (PMI).^19^ PMI was calculated from the lowest docking score pose of each ligand, and as is customary, the normalized PMI coordinates were plotted on a triangular graph. In these graphs, the vertices are associated with rod-like, disc-like, and sphere-like shapes.

## RESULTS AND DISCUSSION

### SampleDock yields target-specific libraries

In this study, we sought to generate a set of target-specific libraries (Figure 1a) of five highly structured RNAs (Figure 1b) to interrogate aspects of the chemical space spanned. Such analysis could help uncover critical features that distinguish the region of chemical space that tend to be compatible with typical RNA-pockets from those of proteins. Therefore, for the FMN Riboswitch, IRES, PreQ_1_, Ribosomal A-Site, and SAM Riboswitch RNAs, we used the SampleDock approach to generate in silico libraries that were compatible with known ligand binding sites on their surface. Briefly, SampleDock combines a pre-trained AI generative model with molecular docking calculations to sample chemical space conditioned on the properties of a specific site on a specific target. Thus, we used to design five target-specific libraries, each targeting a single site on each RNA. In the case of the FMN Riboswitch, PreQ_1_, Ribosomal A-Site, and SAM riboswitch, the site targeted was the site occupied by the cognate ligand. In the case of IRES RNA, the site we targeted was the site occupied by the known inhibitor (PDB ID: SS0).

**Figure 1:**
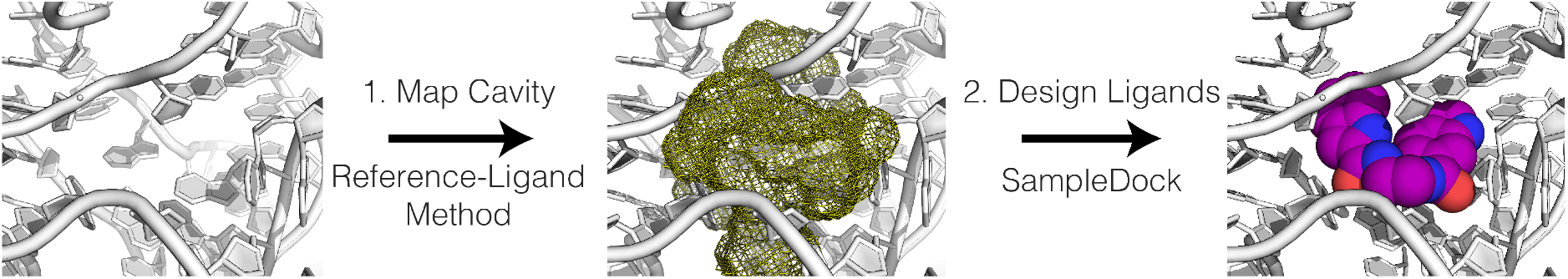
Here we generate and analyze target- and site-specific libraries. Such libraries are generated by first mapping a cavity and second designing ligands that are interact favorably with the nucleotides that line the cavity. Here we map cavities using the reference ligand technique and design ligands using the SampleDock methodology.

First, to confirm that the SampleDock generated libraries were site-specific, we conducted a cross-library docking analysis. In cross-library docking, one docks the library generated for an intended target against sites on other targets. If a library is target- and site-specific, the expectation would be that a library would exhibit elevated docking scores when docked in sites other than the intended, targeted site. Figure 2 shows the violin plots of the docking score distributions from the cross-library docking analysis. For each of the five RNAs we studied, we found that the SampleDock library that was generated by targeting the known ligand-binding site of that RNA (Figure 1a) exhibited significantly lower docking scores than the molecules in the SampleDock libraries that were generated by targeting the known ligand binding sites of that other four RNAs. For instance, the library generated within the FMN binding pocket on the FMN riboswitch had a mean docking score of −32.32 kcal/mol compared with −11.17 kcal/mol for the molecules in the libraries of the other four RNA (Figure 2a). Collectively, the results confirm that the SampleDock framework was able to generate libraries that were specific to a particular site on a given target.

**Figure 2:**
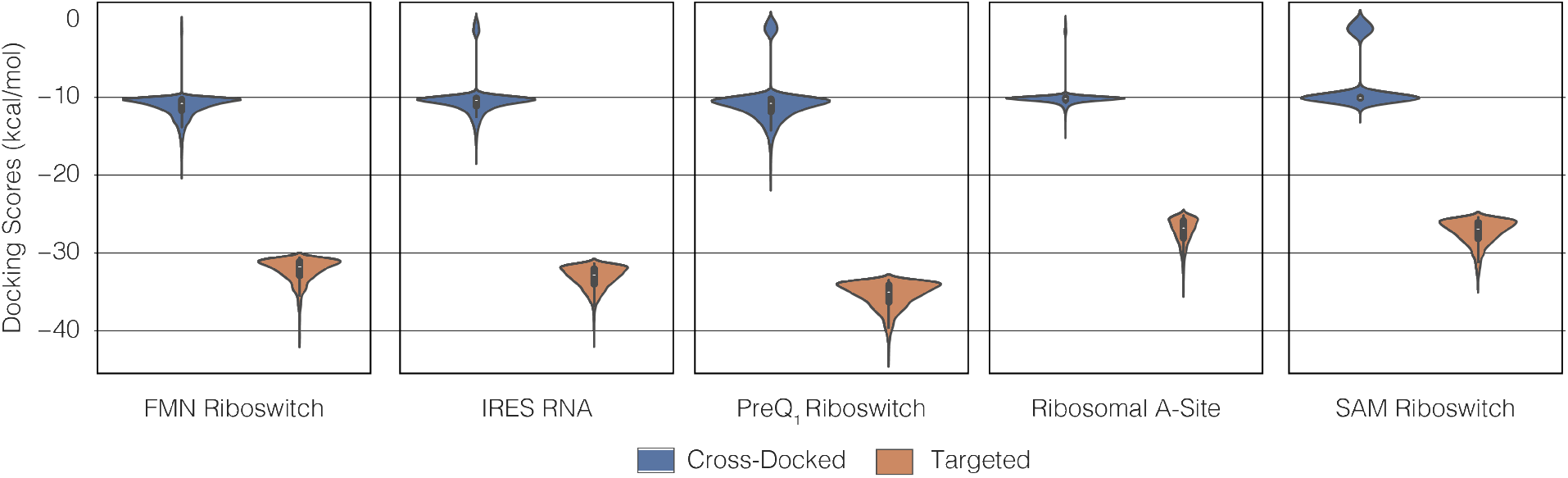
Boxplots showing the distribution of the 20 physiocochemical properties listed in Table S1. Truman, annotate each boxplot with relevant values from the R-BIND paper.

### The distribution of key physicochemical properties within our SampleDock library resemble the distributions in the R-BIND

Satisfied that our libraries were target-specific, we next set out to directly quantify the distribution of several physicochemical properties with the structure-aware libraries. In Table S1 we list the set of properties we monitored. We chose these properties because they are frequently used to describe the drug-likeness of ligands. Furthermore, these properties have recently been used within a cheminformatics analysis to compare the chemical space spanned known bioactive RNA binders to those spanned by bioactive protein binders.

For this analysis, we combined the SampleDock libraries of the five targets. We then compared the distributions of the 18 physicochemical properties across the combined library to the distributions within the R-BIND, the most carefully curated RNA binding database currently available. With few exceptions, and when compared with distributions across the FDA drugs, we observed a high degree of overlap between the SampleDock and R-BIND distributions (Figure 3). Thus, despite being explicitly structure-based, our results are consistent with the conclusion, based on the cheminformatics analysis of the R-BIND, that RNA binders molecules are likely to occupy privileged regions of chemical space relative to known protein targeting molecules.

**Figure 3:**
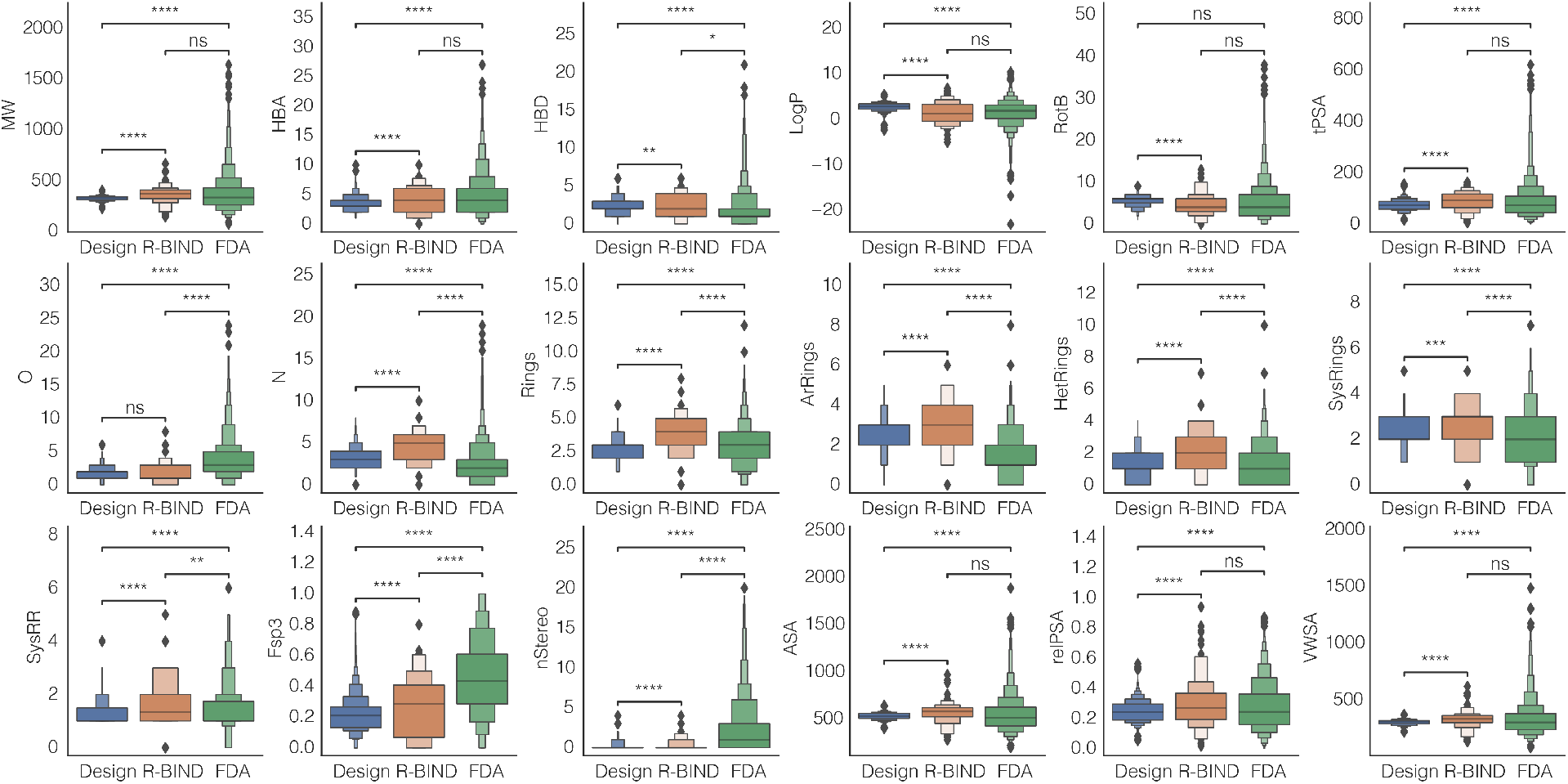
Comparison between SampleDock libraries and R-BIND. Letter-value plots compare distribution of 18 physicochemical properties across the combined SampleDock libraries of the five targets we studied to those in R-BIND.

### Principal moments of inertia (PMI) analysis of SampleDock also suggest that RNA binders may tend to be more rod-like

Using principal moments of inertia (PMI) analysis, Hargrove and coworkers recently discovered that small molecules in their R-BIND tend to be rod-like.^14,17^ However, the PMI analysis is entirely ligand-based and thus agnostic to the biophysical details of the RNA pocket to which these ligands bind. We, therefore, also carried out PMI analysis on the SampleDock libraries, using for the analysis the docked poses of molecules in their respective pockets. Interestingly, we found that molecules in the library also tend to be rod-like (Figure 4), supporting the notion that the tendency towards rod-likeness observed in the R-BIND might be partly linked to the constraints associated with the ligand-binding pockets. We note, however, that the degree of rod-likeness varied with the specific site targeted, with the rod-like character being most pronounced for the library targeting the preQ_1_ riboswitch pocket and least pronounced for the library targeting the FMN riboswitch pocket.

**Figure 4:**
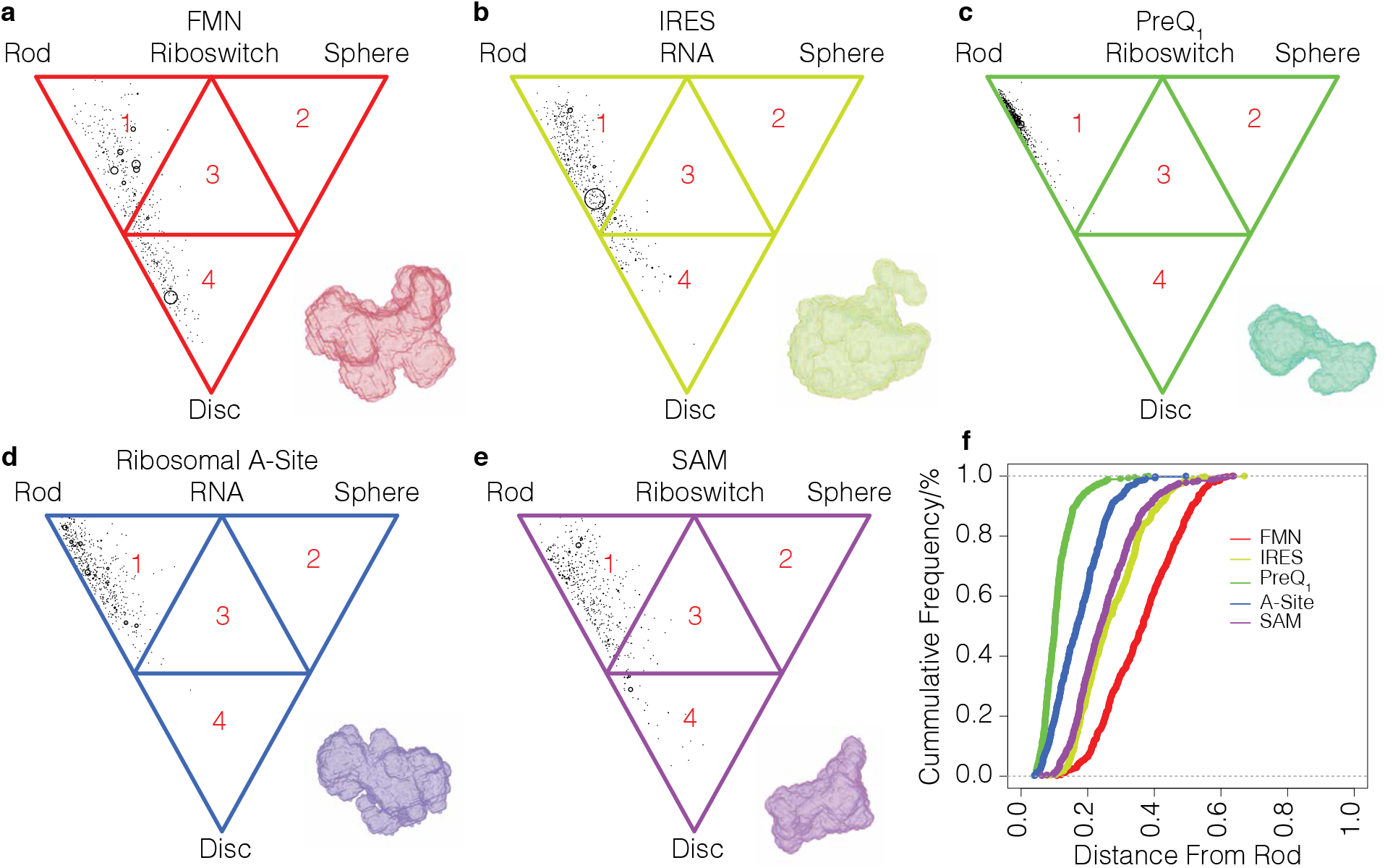
Principal moments of inertia (PMI) triangular plots for the five target-specific RNA libraries (Table 1).

### Nitrogen-containing ring systems are frequently observed in our libraries

Finally, we asked what were the most frequently observed ring systems in the molecules within our virtual libraries. In Figure 5a, we show the 40 ring systems that appeared at least ten times in our combined virtual library; we show all 108 ring systems with frequencies greater in Figure S1. Notably, more than 75% of the rings are nitrogen-containing. To assess whether these rings might represent chemical building blocks that are likely to be found in RNA ligands, we analyzed a collection of 69 small-molecule fragments that were recently validated to bind to 15 distinct RNA elements in the SARS-CoV-2 genome.^15^ Of the 108 ring systems, 32 were present in the 69 SARS-CoV-2 RNA binders. Although, extensive analysis and testing will be required to identify all of the chemical building blocks of RNA-specific ligands, our results provide a preliminary indication that the rings systems frequently observed in our virtual libraries include some of the chemical building blocks of actual RNA binders.

**Figure 5:**
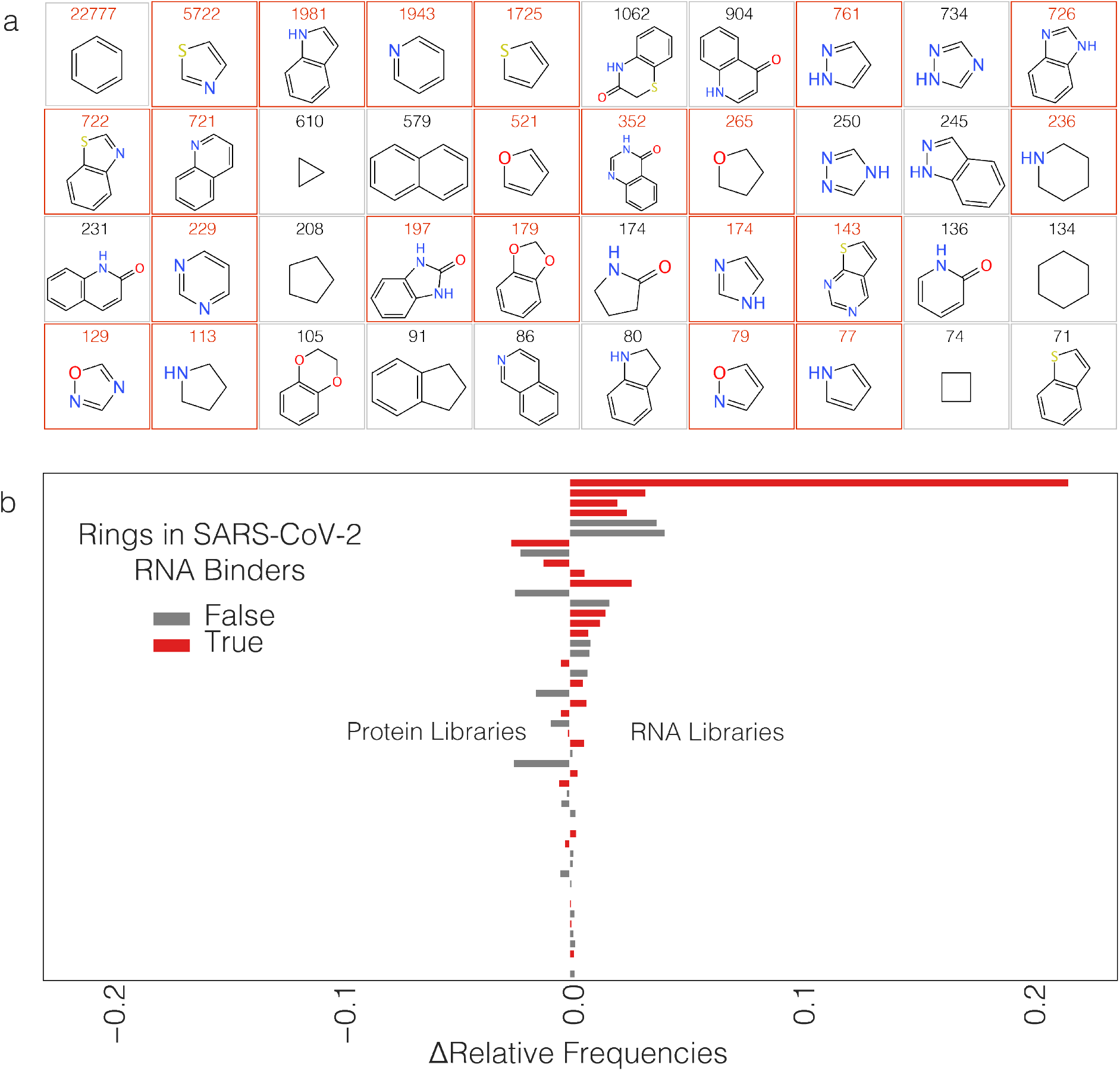
(a) Top 40 ring systems found in our combined virtual library. The ring systems are ranked based on their frequency, which we indicated above the image of each ring system. Here we show ring systems observed in at least ten unique molecules. Rings with red boxes are those found recently reported library of molecules that were validated to bind to various RNA elements within the SARS-CoV-2 genome. ^15^ (b) Relative frequency of top 40 ring systems found in our combined virtual library. For comparison, the relative frequency in a combined library targeting CDK2 and M^Pro^. Highlighted in red are ring systems among the top 40 that are present in validated binders of various RNA elements within the SARS-CoV-2 genome. ^15^

## CONCLUSIONS

In this study, we used a structure-based technique to computationally explore the RNA-specific regions of chemical space. Our strategy was to first generate virtual libraries that were specific for the known ligand binding sites in five highly-structured RNAs and then analyze these libraries.

First, using cross-docking analysis, we found that the libraries we generated were target-specific from a docking standpoint. That is, a library generated for a given target tended to exhibit elevated (less favorable) docking scores when docked to sites other than the one targeted. Second, the distribution of many critical physicochemical properties across our target-specific libraries overlaps with the corresponding distributions in R-BIND. The R-BIND is a curated collection of known bioactive RNA binders. Previous analysis on the R-BIND suggests that RNA targeting small molecules spans a distinct region of chemical space compared with those known to target the proteins. Third, like in the R-BIND, distributions of O, N, Rings, ArRings, HetRings, SysRings, and Fsp3 in our libraries exhibited small p-values relative FDA approved. And fourth, our libraries share a characteristic feature with molecules in the R-BIND, namely a tendency to be rod-like. We conclude by highlighting a key limitation of our current study, namely, that due to computational cost, we generated target-specific libraries for five targets. Thus the comparison between our libraries and the R-BIND must be framed by the small sample size. Our results, however, strongly suggest that a more extensive analysis of the known RNA “pocketome” might yield valuable insights about the regions of space that span those that include RNA-specific binders. Furthermore, compared with a slower and more costly wet-lab-then-chemoinformatics approach, such a computationally-driven approach is the most efficient approach we can currently adopt to generate mental models about the molecular recognition features of RNA and how they differ from proteins.

## ACKNOWLEDGMENT

We thank the members of the Frank lab for many valuable discussions about this work. The authors were funded by startup funds and a grant from the Michigan Institute for Computational Science and Discovery (MICDE) at the University of Michigan (Z.X. and A.T.F).

## ADDITIONAL NOTES

### SUPPORTING INFORMATION DESCRIPTION

This material is available free of charge via the Internet at http://pubs.acs.org.

## DATA AND SOFTWARE AVAILABILITY

SampleDock can be accessed as a web service from SMALTR at https://smaltr.org or install locally following the instructions in https://github.com/atfrank/SampleDock.

## SUPPORTING INFORMATION

**Table S1:** Chemical Properties Used to Characterize and Compare Libraries. ^18^

| Parameter | Description |
| --- | --- |
| MW | Molecular Weight |
| HBA | Number of Hydrogen Bond Acceptors |
| HBD | Number of Hydrogen Bond Donors |
| LogP | n-Octanol/Water Partition Coefficient |
| RotB | Number of Rotatable Bonds |
| tPSA | Topological Polar Surface Area |
| O | Number of Nitrogen Atoms |
| N | Number of Oxygen Atoms |
| Rings | Number of Rings |
| ArRings | Number of Aromatic Rings |
| HetRings | Number of Heteroatom-containing Rings |
| SysRings | Number of Ring Systems |
| SysRR | Ring Complexity |
| Fsp3 | Fraction of sp<sup>3</sup> Hybridized Carbons |
| nStereo | Number of Stereocenters |
| ASA | Accessible Surface Area |
| relPSA | Relative Polar Surface Area |
| VWSA | Van der Waals Surface Area |

**Figure S1:**
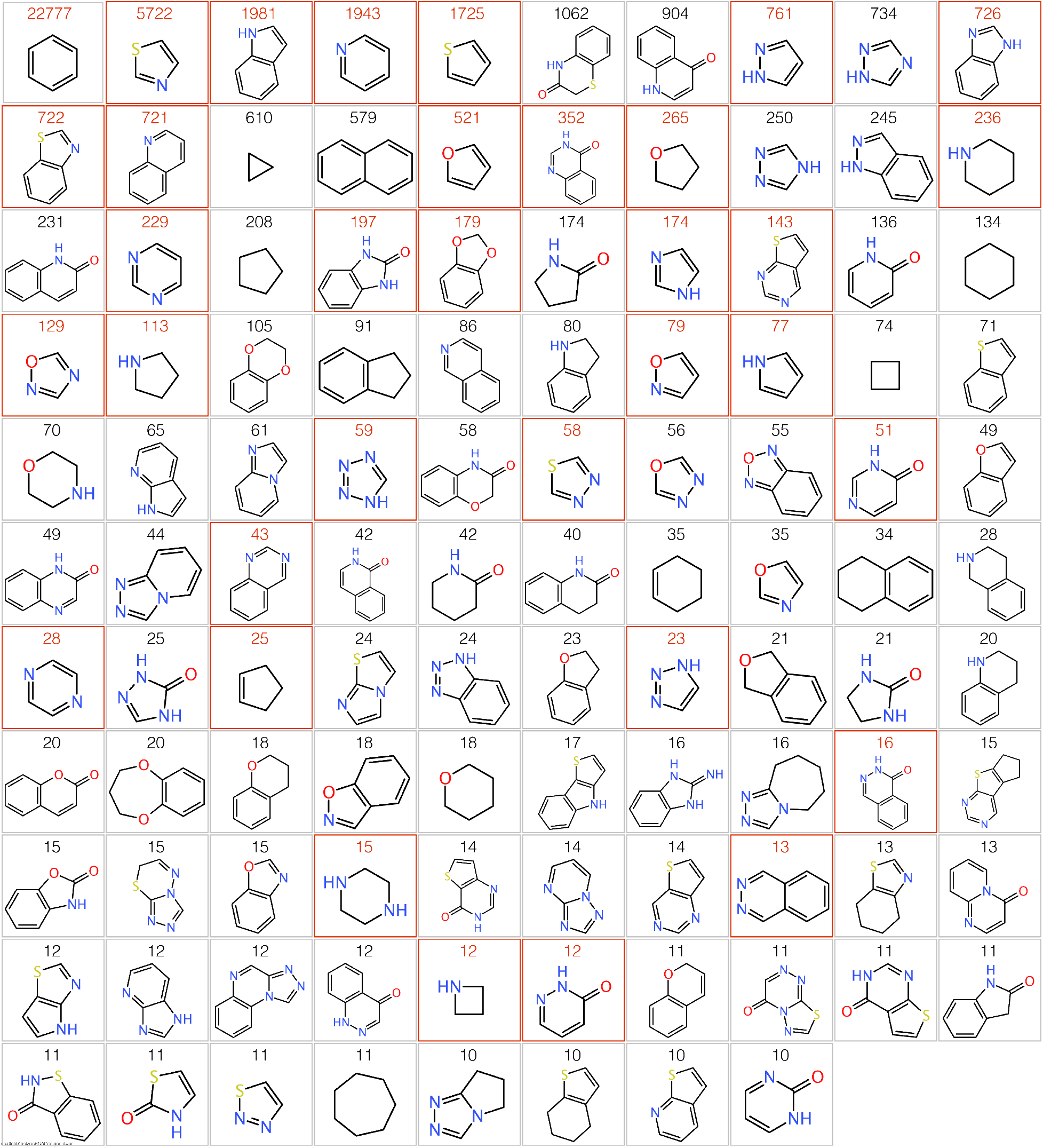
Ring systems found in our combined virtual library. The ring systems are ranked based on their frequency, which we indicate above the image of each ring system. We show ring systems that we observe in at least ten unique molecules. Rings with red boxes are those found recently reported library of molecules that were validated to bind to various RNA elements within the SARS-CoV-2 genome.^15^

